# Comparative transcriptome analysis of mule uncovered several heterosis-related genes for improving muscular endurance and cognition

**DOI:** 10.1101/823047

**Authors:** Shan Gao

## Abstract

Heterosis has been widely exploited in animal and plant breeding to enhance the productive traits of hybrid progeny of two breeds or two species. Although, there were multiple models for explaining the hybrid vigor, such as dominance and over-dominance hypothesis, its underlying molecular genetic mechanisms remain equivocal. The aim of this study is through comparing the different expression genes (DEGs) and different alternative splicing (DAS) genes to explore the mechanism of heterosis. Here, we performed a genome-wide gene expression and alternative splicing analysis of two heterotic crosses between donkey and horse in three tissues. The results showed that the DAS genes influenced the heterosis-related phenotypes in a unique than DEGs and about 10% DEGs are DAS genes. In addition, over 69.7% DEGs and 87.2% DAS genes showed over-dominance or dominance, respectively. Furthermore, the “Muscle Contraction” and “Neuronal System” pathways were significantly enriched both for the DEGs and DAS genes in muscle. *TNNC2* and *RYR1* genes may contribute to mule’s great endurance while *GRIA2* and *GRIN1* genes may be related with mule’s cognition. Together, these DEGs and DAS genes provide the candidates for future studies of the genetic and molecular mechanism of heterosis in mule.

## Introduction

Heterosis refers to the phenomenon that hybrids exhibit superior performance such as stress resistance, growth rate, and biomass production relative to either of their parents (Birchler et al., 2003; Hochholdinger and Hoecker, 2007; Zhang et al., 2008), and heterosis has been widely used in increasing the quantity of crops and livestock breeding. Three classic genetic hypotheses: dominance (Davenport, 1908; Bruce, 1910), over-dominance (Shull, 1908), and epistasis (Yu et al., 1997) had been built to explain the heterosis. The dominance model proposes that the distinct sets of deleterious recessive alleles undergo a genome-wide complementation in hybrids. In addition, the over-dominance model states that the intra locus allelic interactions at one or more heterozygous genes leads to increased vigor. Unlike two models as above mentioned are based on a single locus to explain heterosis, the epistasis is defined as the interaction of the dominant alleles at different loci from the parents, i.e. the trait of a certain gene affected by one or more other genes. Although the dominance and over-dominance hypotheses can be reflected on the expression profile of genes decently, it’s still difficult to explain the heterosis integrally. With the development of functional genomics and the improvement of RNA sequencing (RNA-seq) technology, heterosis in cattle (Bunning et al., 2019), mice (Ebato et al., 1983), wheat (Sun et al., 2004; Li et al., 2014), rice (Zhang et al., 2008; Wei et al., 2009), corn (Swanson-Wagner et al., 2006), tomato (Krieger et al., 2010) were investigated at the transcription level from the point of gene expression.

Alternative splicing is a process in which a gene transcribes from the same precursor mRNA and then undergoes a splicing process to produce two or more mature mRNAs, ultimately producing different proteins(Black, 2003). Furthermore, the varied proportion of different splicing isoforms also influences the phenotype, which may also contribute heterosis. For example, two transcriptome isoforms of the *Titin* gene expressed in the different developmental stage of heart: the long isoform *N2BA* of the *Titin* gene predominantly during the development, while the shorter isoform *N2B* of one’s gene dominates in adults (Krüger and Linke, 2011; Guo et al., 2012; 2012; Li et al., 2013). If the *N2BA* isoform mainly expressed in the heart of adults, it can cause heart disease (Brauch et al., 2009; Haas et al., 2015). However, analysis on alternative splicing for hybrids vigor are rarely reported, so detecting the genes with differentially alternative splicing (DAS) events between hybrids and either of their parents is essential.

Mule, the hybird of male donkeys and female horses, differed from the hinny and represented as an example of hybrid vigor, exhibiting better and stronger muscle endurance and cognition than either of their parents and hinny. In addition, the mule exhibited obviously difference in height, body size, hair etc. either of their parents and hinny. (Figure 1A) (Proops et al., 2009; Osthaus et al., 2013; Renaud et al., 2018). In this study, we collected the muscle, brain, skin from mule and their parents and hinny and detected Differentially Expressed Genes (DEGs) and different alternative splicing (DAS) in order to explore the genetic and molecular mechanisms of heterosis-related phenotype in mule. Our results showed that the mule is different from the hinny and both of their parents at the expression level of gene and differential PSI (percent spliced-in) in alternative splicing events. The detection of DEGs and genes with DAS events provided the heterosis-related candidate genes which may be an important contributor to heterosis.

**Figure 1:**
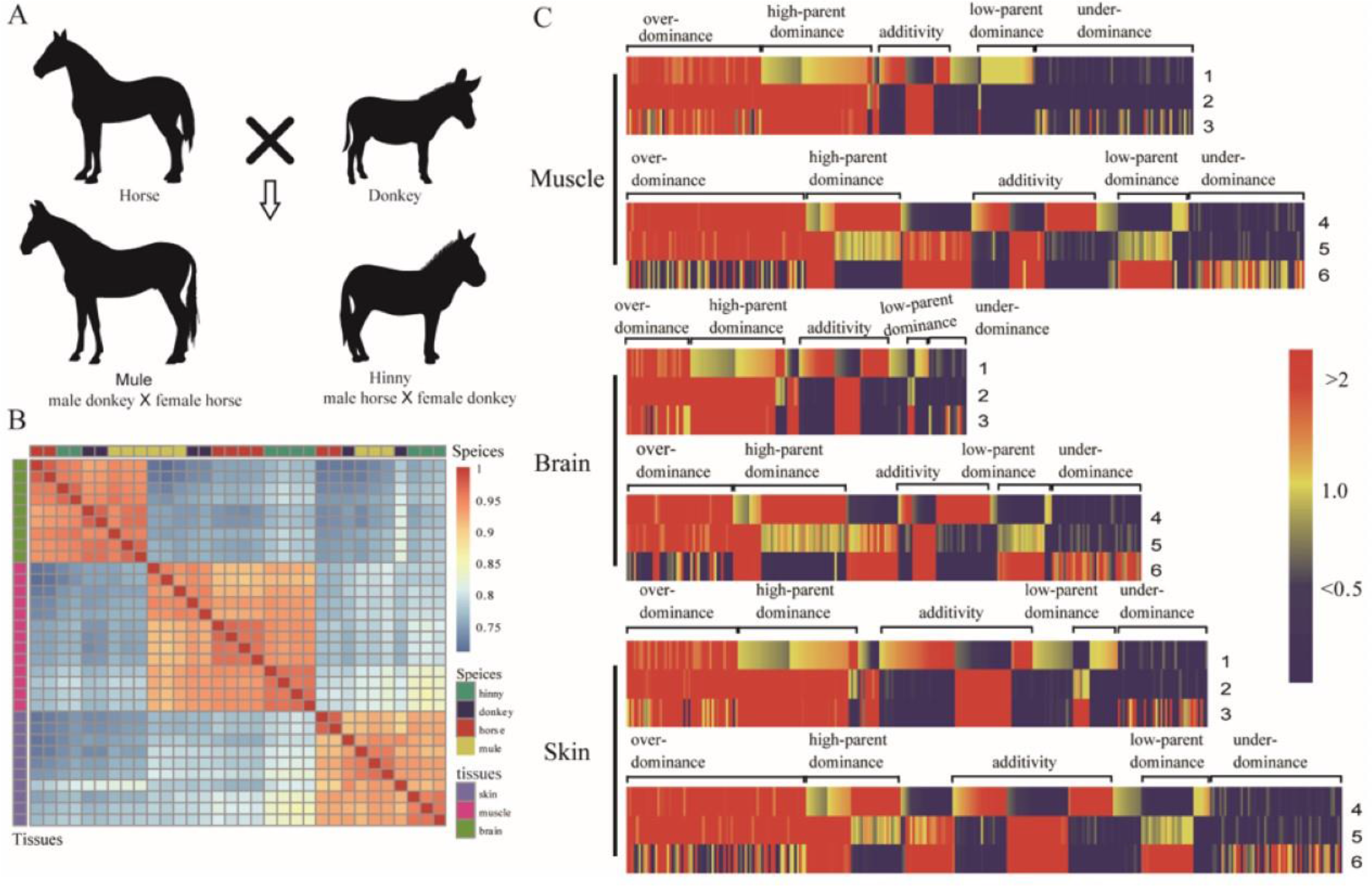
Differentially expressed genes in the tissues of muscle, brain and skin from mule, hinny and their parents. (A) The corss between horse and donkey. Left panel is mule and its parents (male donkey × female horse), right panel is hinny and its parents (male horse × female donkey). (B) The Pearson’s correlation of the expression profile of genes in the tissues of brain, muscle and skin from mule, hinny and their parents is showed on the top and the clustered results of gene expression among species are shown on the top, while the clustered results of gene expression in these tissues of species are shown on the side. (C) Hierarchical clustering display of differential expressed genes with the ratio of intensity according to expression patterns of genes among mule, hinny and their parents. The color scale is shown at the top and mode of gene action is in the side. Lane 1 represents intensity ratios of mule to horse; Lane 2, intensity ratios of mule to donkey; Lane 3, intensity ratios of horse to donkey. Lane 4 represents intensity ratios of hinny to horse; Lane 5, intensity ratios of hinny to donkey; Lane 6, intensity ratios of horse to donkey.

## Materials and Methods

### Animal materials and data collection

The samples of donkeys, hinnies and mules are as the same as the samples in the Allele-specific Expression (ASE) program(PRJNA387435) conducted by previous research (Wang et al., 2019). The RNA samples used for generating Pac-Bio long reads were extracted from the same mule (number:M2M) as in PRJNA387435. The sequencing raw data has been upload to NCBI (accession number: PRJNA560325). The RNA-Seq data of horse were collected from NCBI (accession number: PRJNA339185 and PRJEB26787). The study was approved by the Institutional Animal Care and Use Committee of Northwest A&F University (Permit Number: NWAFAC1019).

### RNA preparation and single-molecule sequencing

Total RNA was extracted from the muscle tissue of the mule using Trizol reagent according to the protocol of manufacturer. The concentration was measured by NanoDrop 2000 spectrophotometer (Thermo Scientific, USA) and the quality was estimated by Agilent Bioanalyzer 2100. Five different size fragment sequencing libraries (1-2kb, 3*2-3kb, >3kb) were constructed according to the guide for preparing SMRT bell template for sequencing on the PacBio RS platform.

### RNA reads trimming and alignment

The RNA-seq raw reads were first removed adapter sequences. Then, the reads were trimmed using Trimmomatic (v0.33) (Bolger et al., 2014) and the length of reads lengther than 70 nucleotides were retained as high quality clean data. The clean reads of high quality from mule/hinny were aligned to the horse reference genome (Equus caballus: EquCab2.0) by STAR (v 2.5.1a) (Dobin et al., 2013).

### Differential expression analysis

We used Stringtie (v 1.3.4d) (Pertea et al., 2015; Pertea et al., 2016) and the attached python script prepDE.py (http://ccb.jhu.edu/software/stringtie/index.shtml?t=manual) to quantify the reads counts and calculated DEGs by DESeq2 R package (v 1.10.1) (Love et al., 2014). The corrected *P*-values (Benjamin-Hochberg test) less than 0.05 were deemed to significant DEGs.

### Functional enrichments analysis

KEGG pathway enrichment analysis were carried out by the online software kobas (v3.0) (http://kobas.cbi.pku.edu.cn/) (Xie et al., 2011). The protein sequences of genes with DAS and DEGs were submitted to human database and hypergeometric test/Fisher’s exact test was used to calculate the corrected *P*-values.

### Analysis of DAS genes

Replicate Multivariate Analysis of Transcript Splicing (rMATs, v3.2.5) (Shen et al., 2014) was utilized for identification and comparison of alternative splicing events of genes, including Exon skipping (SE), mutually exclusive exons (MXE), alternative 5’ splice site (A5SS), alternative 3’ splice site (A3SS) and retained intron (RI). The significance testing of the results from rMATs software using a likelihood-ratio method to calculate the P value based on the differential ψ values also named “percent spliced-in” (PSI). The splicing events were assessed with the rigorous statistical criteria (|Δψ|> 10% and FDR ≤ 0.05) which quantified DAS.

### PacBio full length transcripts analysis

The PacBio RS SMRT sequence reads were first analysis on the Pacific Biosciences’ SMRT analysis software (v2.3.0) to get reads of insert (ROI). Then, the ROIs were aligned to the correspond reference genome using GMAP (v 2015-12-31) (Wu and Watanabe, 2005). The high error rate of long reads were corrected by TAPIS (https://bitbucket.org/comp_bio/tapis). Then the differentially alternative splicing events were visualized using R package Gviz (v 1.18.1)(Hahne and Ivanek, 2016).

## Result

### Overview of the gene expression level in the tissues of brain, muscle and skin from mule

To explore the effects of differentially gene expression on heterosis in mule, the RNA-seqs were performed between the hybrids and either of their parents. After removing the adaptor and low-quality reads, over 427, 601 and 482 million clean reads were obtained from the tissues of brain, muscle and skin, respectively. The average mapping ratios in the above three tissues were 91.50%, 90.22% and 91.81%, respectively. The expression profile of 24,331 DEGs identified among those three tissues with log2 transformed > 1 and normalized by DEseq2. The correlation analysis of gene expression was implemented, it indicated that all samples were clustered by tissues firstly, and then the same species tended to be cluster together (Figure 1B). The similar pattern of results were shown from Principal Component Analysis (PCA) (Figure S1).

### Analysis of DEGs between the hybrids and either of their parents

A total of 3,689, 7,044 and 6,637 DEGs (adjust-*p* < 0.05) were identified from the tissues of brain, muscle and skin between mule and horse, respectively, in which 2,162 (58.6%), 3,442(48.9%) and 3,560 (53.6%) DEGs exhibited up-regulated in the above three tissues, while 3,602 (51.1%), 1,527 (41.4%) and 3,077 (46.4%) DEGs exhibited down-regulated, respectively. On the other side, 952, 275 and 1,326 DEGs (adjust-*p*-value < 0.05) were identified from the tissues of brain, muscle and skin between mule and donkey, respectively. 611 (64.2%), 202 (73.5%) and 606 (45.7%) DEGs showed up-regulated in the above three tissues, while 341 (35.8%), 73 (26.5%) and 720 (54.3%) DEGs showed down-regulated, respectively. Similarly, the DEGs between hinny and either of their parents were also detected (Table 1 and Supplementary Figure 2). Then we divided DEGs identified between mule and horse into four patterns according to the classic genetic hypotheses on heterosis based on their deviation from the mid-parent prediction, including high-parent dominance genes, low-parent dominance genes, over-dominance genes, and under-dominance genes and found 7,196 (26.5%), 4,334 (16.0%), 7,367 (27.1%) DEGs (adjust-*p* - value< 0.05) (Figure S2) were identified in the tissues of muscle, brain and skin between mule and either of their parents, respectively. Among these genes, 6,762 (94.0%), 3,709 (85.6%) and 6,144 (83.4%) DEGs in the tissues of muscle, brain and skin showed non-additive pattern, respectively. While the DEGs in the pattern of additive were classified as over-dominace and under-dominance following the hypothesis of over-dominace and those DEGs in the same pattern were classified as high-parent dominace and low-parent dominace following the hypothesis of dominace. The majority of DEGs showed over-dominance in the samples we collected, such as followings: 52.3%, 30.1% and 34.4% DEGs in the above three tissues showed actions of over-dominance and under-dominance modes, while 33.8%, 39.6% and 35.3% DEGs exhibited high-parent dominance and low-parent dominance modes, respectively (Table 2). We also found the expression patterns of gene in the tissues of brain, muscle and skin from mule are more closely related to the donkey than the horse based on the hierarchical clustering algorithm (Figure 1C), however, the expression patterns of gene in those tissues from hinny are diversely from either of their parents. These results suggested that the gene expression patterns in hybrids are close related to either paternal or maternal.

**Table 1:** The distribution of differentially expressed genes across the tissues of muscle, brain and skin in the mules, hinnies and their parents.

| Tissues | Gene type | Mules vs horses | Mules vs donkeys | Hinnies vs horses | Hinnies vs horses |
| --- | --- | --- | --- | --- | --- |
| Muscle | Up-regulated | 3,602 | 73 | 2,523 | 2,442 |
|  | Down-regulated | 3,443 | 203 | 2,811 | 3,029 |
| Brain | Up-regulated | 1,527 | 341 | 1,054 | 2,308 |
|  | Down-regulated | 2,163 | 612 | 1,224 | 2,896 |
| Skin | Up-regulated | 3,077 | 720 | 2,899 | 2,573 |
|  | Down-regulated | 3,561 | 607 | 3,312 | 2,646 |

**Table 2:** The classification of differentially expressed genes following the expression patterns of genes. Hybirds represents mule or hinny; P, paternal lines represent male horse or male donkey; M, maternal lines represent female horse or female donkey. Additivity, mule or hinny = 1/2(horse+donkey); non-additivity, mule or hinny > 1/2(horse+donkey) or mule or hinny < 1/2(horse+donkey). High-parent dominance, mule or hinny = P > M or mule or hinny = M > P; low-parent dominance, mule or hinny = P < M or mule or hinny = M < P; over-dominance, mule or hinny > P and mule or hinny > M; under-dominance, mule or hinny < P and mule or hinny < M

| Tissues | Hybrids | Non-addictive | Over-dominance | High-parent dominance | Addictive | Low-parent Dominance | Under-dominance |
| --- | --- | --- | --- | --- | --- | --- | --- |
| Muscle | Mule | 561 | 1,713 | 1,397 | 434 | 1,040 | 2,051 |
|  | Hinny | 1,135 | 2,281 | 1,207 | 1,371 | 1,160 | 1,459 |
| Brain | Mule | 688 | 811 | 1,212 | 625 | 503 | 495 |
|  | Hinny | 992 | 1,369 | 1,431 | 826 | 792 | 1,148 |
| Skin | Mule | 1,008 | 1,394 | 1,532 | 1,223 | 1,069 | 1,141 |
|  | Hinny | 1,363 | 2,294 | 1,189 | 1,343 | 1,237 | 1,665 |

To investigate the possible biological function of DEGs in hybirds, pathway enrichment analysis of the DEGs were conducted by Kobas. (Table S1, and Figure S3, S4, S5). The DEGs identified from the tissues of muscle, brain and skin in hybirds were significantly enriched in “Muscle contraction”, “Neuronal System” and “DNA Repair” pathways, respectively.

### Identification and Characterization of differential alternative splicing events

A total of 2,241, 2,657 and 1,973 AS events were identified from 1,240, 1,674 and 1,330 genes in the tissues of muscle, brain and skin from mule, respectively (Figure S6). The samples were clustered by tissue type firstly based on the gene alternative splicing (PSI value) and then the same species were tended to cluster together (Figure 2A), which is consisted with the results of gene expression. Similarly, the genes with DAS in hinny were also detected in the three tissues as mentioned above (Table 3). The genes with DAS events were also divided into four categories based on the same standards as the expression profiles of genes and the inclusion level of the alternative splicing events: the high-parent dominance, low-parent dominance, over-dominance, and under-dominance. (Figure 2B and Table4). As depicted in Table 4, the highest proportion of genes with alternative splicing events in the above three tissues showed high-parent dominance, followed by low-parent dominance, Under-dominance and Over-dominance. The results of pathway enrichment analysis on genes with DAS in mule showed that “Muscle contraction”, “Neuronal System” pathways were significantly enriched in the tissues of muscle and brain, respectively (Table S2, and Figure S7, S8, S9). Genes exhibited over-dominance were significantly enriched in the pathways of Vasopressin synthesis, Circadian rhythum, while genes showed dominance were significantly enriched in the pathways of Striated Muscle Contraction, Muscle contraction (Figure 2C, 2D).

**Figure 2:**
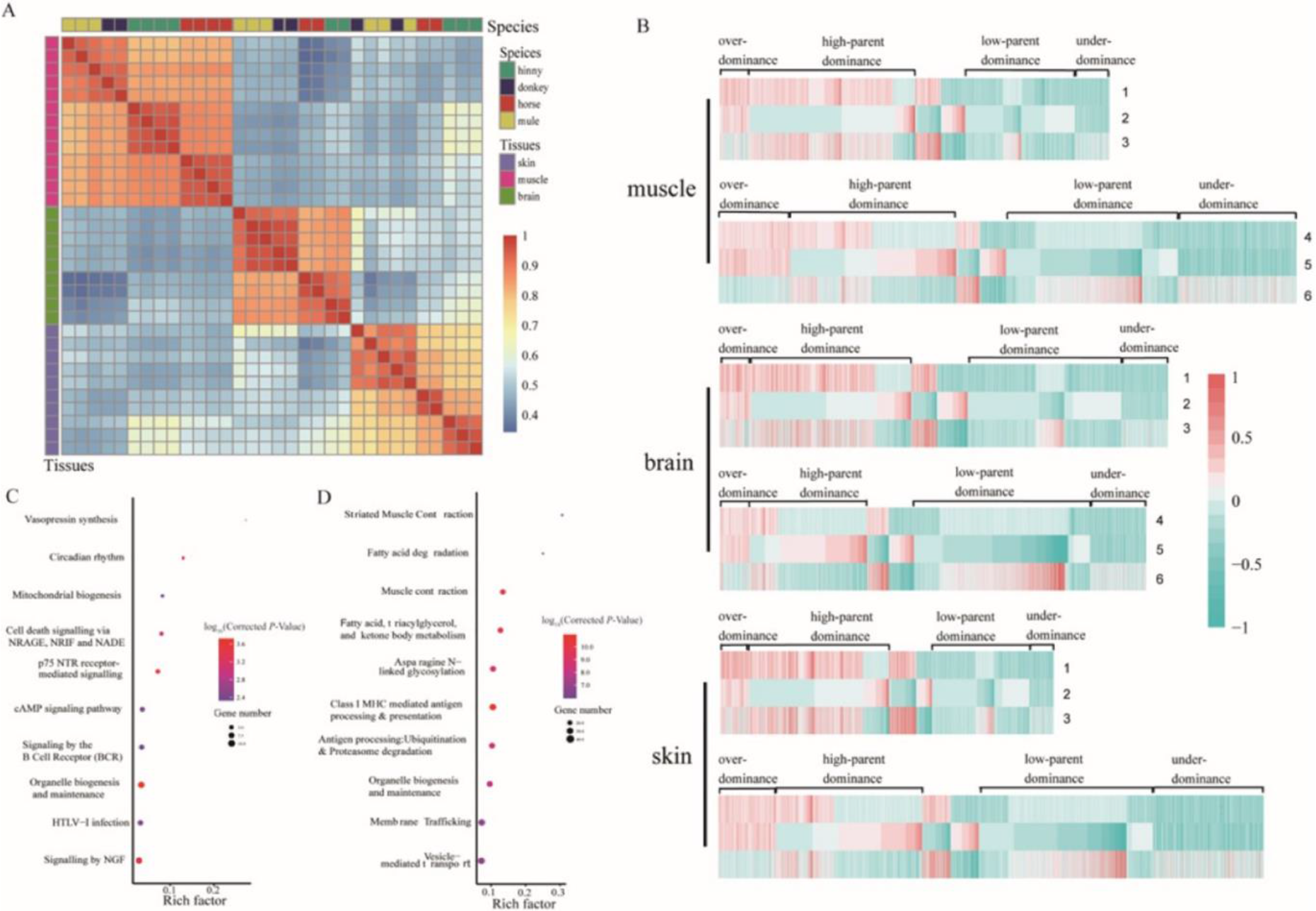
The genes with differentially alternative splicing events in the tissues of muscle, brain and skin from mule, hinny and their hybirds. (A) The Pearson’s correlation coefficient of the PSI (percent spliced-in) of genes with DAS in the tissues of muscle, brain and skin from mule, hinny and their parents. the clustered results of genes with differential alternative splicing events among species are shown on the top, while the clustered results genes with differential alternative splicing events among species are shown on the side. (B) Hierarchical clustering display of DAS genes with the ratio of intensity according to expression patterns among horse, donkey mule and hinny. The color scale is shown at the top and the mode of gene action is shown in the side. Lane 1 represents intensity ratios of mule to horse; Lane 2, intensity ratios of mule to donkey; Lane 3, intensity ratios of horse to donkey. Lane 4 represents intensity ratios of hinny to horse; Lane 5, intensity ratios of hinny to donkey; Lane 6, intensity ratios of horse to donkey. (C) The KEGG pathways enriched by the genes that following the hypothesis of over-dominance. (D) The KEGG pathways enriched by the genes that following the hypothesis of dominance.

**Table 3:** The distribution of the Genes with differentially alternative splicing events across the tissues of muscle, brain and skin in the mules, hinnies and their parents.

| Tissues | Mules vs horses | Mules vs donkeys | Hinnies vs horses | Hinnies vs horses |
| --- | --- | --- | --- | --- |
| Muscle | 1377 | 593 | 656 | 1319 |
| Brain | 1124 | 280 | 1264 | 1330 |
| Skin | 1128 | 378 | 1121 | 1570 |

**Table 4:** The classification of genes with differentially splicing events following the expression patterns of genes. Hybirds represents mule or hinny; P, paternal lines represent male horse or male donkey; M, maternal lines represent female horse or female donkey. Additivity, mule or hinny = 1/2(horse+donkey); non-additivity, mule or hinny > 1/2(horse+donkey) or mule or hinny < 1/2(horse+donkey). High-parent dominance, mule or hinny = P > M or mule or hinny = M > P; low-parent dominance, mule or hinny = P < M or mule or hinny = M < P; over-dominance, mule or hinny > P and mule or hinny > M; under-dominance, mule or hinny < P and mule or hinny < M

| Tissues | Hybrids | Over-dominance | High-parent dominance | Addictive | Low-parent Dominance | Under-dominance |
| --- | --- | --- | --- | --- | --- | --- |
| Muscle | Mule | 173 | 950 | 287 | 633 | 198 |
|  | Hinny | 450 | 1037 | 311 | 1075 | 744 |
| Brain | Mule | 189 | 946 | 334 | 910 | 278 |
|  | Hinny | 175 | 695 | 278 | 1043 | 333 |
| Skin | Mule | 171 | 828 | 253 | 578 | 143 |
|  | Hinny | 340 | 866 | 335 | 1033 | 655 |

**Table 5:** The results of genes with differentially alternative splicing events verified by the Pac-Bio long reads.

|  | mule vs horse | mule vs donkey |
| --- | --- | --- |
| Covered DAS genes | 1123 | 279 |
| Verified DAS genes | 768 | 183 |
| ratio (%) | 68 | 66 |

### The DEGs and DAS genes in muscle contraction and Neuronal System pathways

Combining the analysis results of DEGs and genes with DAS, we found 5.8-11.2%, 3.4%-10.9% and 4.0%-12.6% DEGs with DAS from the tissues of muscle, brain and skin between mule and hinny (Figure S10). Furthermore, we also found DEGs and genes with DAS identified from the tissue of muscle were significantly enriched in “Muscle contraction” pathway (Figure 3). A total of 68 genes were mapped in this pathway, including 50 DEGs, and 8 genes (*TNNC2, RYR1, STIM1, CAMK2D, CAMK2B, CACNA1S, DYSF* and *ATP2A1*) of which overlap with 26 DAS genes. 58.0% (29 of 50) DEGs exhibited dominance, while 92.3% (24 of 26) genes with DAS showed over-dominance effects on muscle endurance. The expression level of *TNNC2* gene is mainly expressed in the fast skeletal muscle from the hybirds and horse / donkey, what’s more, the expression level of this gene in mule is twice as much as in horse (Figure 4A). An exon skip event was found in the exon of this gene chr22:34802038-34802136, when the transcriptome isoform of this gene is mainly expressed in the mule. However, *TNNC2* gene is more likely to retain the exon when another transcriptome isoform of this gene is mainly expressed in horse (Figure 4C and Figure S11) and this isoform expressed in the tissue of muscle is more than two times than in horse. In addition, The *RYR1* gene was also been detected in this pathway, it has higher expression level in skeletal muscle than in other tissues (Figure 4B). The *RYR1* gene is in the same condition which contained a SE events in chr10:9571134-9571148 (Figure 4D and Figure S12). In comparison with the expression level of this gene in mule, it is more than twice as much as horse. An exon skip event was also detected in *RYR1* when it is expressed in the muscle tissue of the horse, while the exon tends to remain as it is expressed in mule (Figure 4D). Similarly analysis were conduct on the tissue of brain, a total of 13 DEGs and genes with DAS (*GRIA2*, *GRM5, GRIN1, GABRA1, GABRA2, DLG2, CACNB4, EPB41L5, BZRAP1, TSPAN7, ADCY1, NRXN1, CREM*) were significantly enriched in the Neuronal System pathway. On the other side, 68% DAS genes identified from muscle tissue of mule were verified by the long reads generating from the sequencing platform of the PacBio. The results further illustrated that our analysis of alternative splicing is relative reliable (Figure S13).

**Figure 3:**
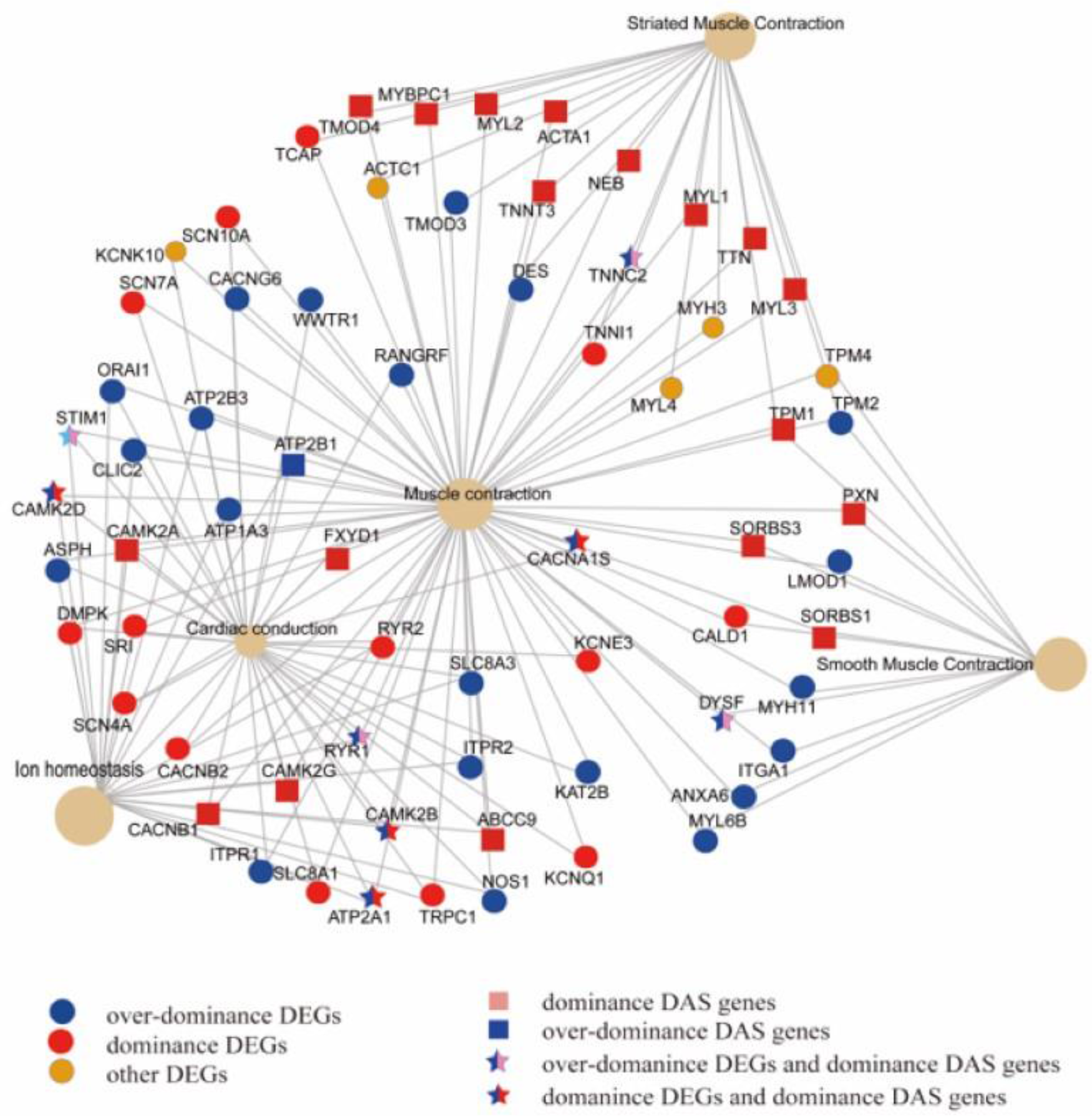
The DEGs and genes with DAS belonging to the models of over-dominance and dominance were enriched in the pathway of muscle contraction.

**Figure 4:**
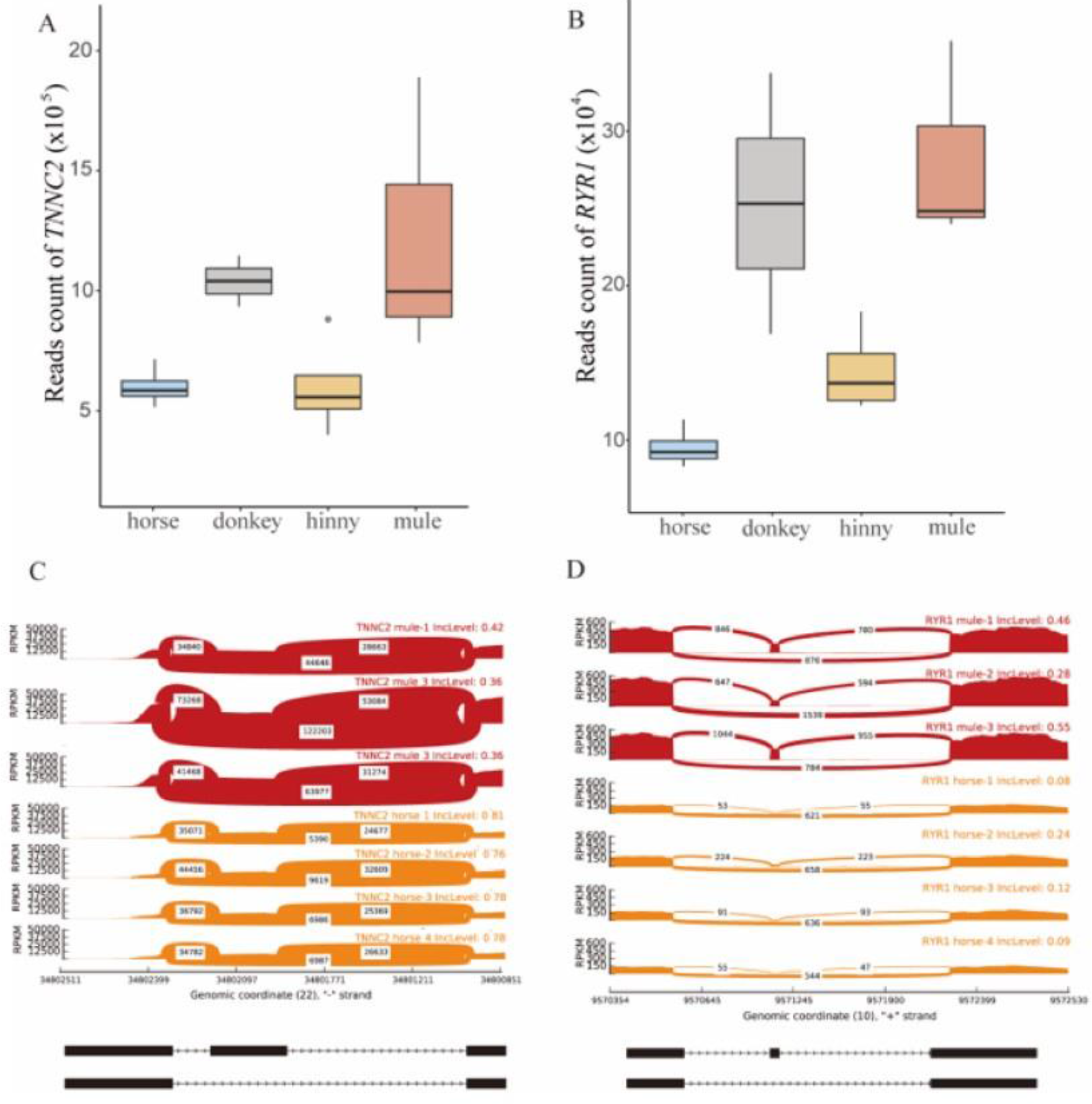
Analysis of *TNNC2* and *RYR1* gene. (A) The distribution of reads mapped on the *TNNC2* gene in the tissue of muscle from mule, hinny and their parents. (B) The distribution of reads mapped on the *RYR1* in the tissue of muscle from mule, hinny and their parents. (C) Sashimi plot of *TNNC2* gene showing DAS in the muscle tissue of mule. (D) Sashimi plot of *RYR1* gene showing DAS in the muscle tissue of mule. Read densities supporting inclusion and exclusion of exons are shown.

## Discussion

The mule is one of the most important conveyance for the people who living in the mountainous area, owing to their outstanding performance in muscular endurance, cognition, growth rate and weight in comparison with either of their parents (Proops et al., 2009; Osthaus et al., 2013; Renaud et al., 2018). However, the genetic and molecular mechanisms of heterosis about mule are unknown. Therefore, we collected the transcriptome data of the tissues of muscle, brain and skin from mule, hinny and their parents and conducted analysis of differential gene expression and genes with differential alternative splicing between hybrids and either of their parents (Guo et al., 2012; Li et al., 2013). From the results of the correlation of gene expression level and alternative splicing events content, it can be see that the divergence between animals can be indicated by the DEGs and DAS genes.

### Comparative analysis between hybrids and either of their parents

We identified a subset of differentially expressed genes in the tissues of brain, muscle and skin between hybrids and either of their parents through the analysis of comparative transcriptome. The correlations of gene expression level and genes with alternative splicing events were first clustered in the same tissue and then clustered in one species, it showed that samples we collected were relative reliable. The gene expression profiles in the tissues of brain, muscle and skin from mule and donkey were clustered together, while the gene expression profiles in the tissue of brain from hinny and horse were clustered together. And the phenomenon extends to genes with alternative splicing events. These results suggested that there are much more difference between mule and hinny at the level of gene expression, although the genetic materials are same between in mule and hinny. Moreover, DEGs were mainly showed over-dominace while the proportion of genes with DAS exhibited dominace is higher than other models. These difference caused by gene expression profiles and alternative splicing events may be the genetic basis of heterosis in the mule.

### Differential alternative-splicing contribute to the heterosis of hybrids

Alternative splicing events are often correlated with the diseases in human when the proportion of isoforms is mis-regulated. For example, if the proportion of distinct isoforms generated from *Titin* gene is abnormal, it can lead to heart disease in human adults (Guo et al., 2012; Li et al., 2013). In this study, we performed a genome-wide analysis of genes with DAS at the transcriptional level. We first identified genes with DAS in different tissues from the same specie through comparative transcriptome analysis, and then compared those genes identified from hybirds and either of their parents by establishing a correlation coefficient matrix among different tissues according to the Pearson’s correlation coefficient. Moreover, the correlation matrix of DAS genes is more relevant in hybirds and either of their parents. These results suggested that differential splicing events between hybirds and either of their parents can reflect the heterosis of hybrids.

### Muscle contraction and Neuronal System pathways in heterosis

Combining the results of KEGG enrichments analysis on DEGs and genes with DAS, we found Muscle contraction and Neuronal System pathways were significantly enriched simultaneously. A total of 8 and 13 DEGS and genes with DAS were significantly enriched in Muscle contraction and Neuronal System pathways, respectively. According to the expression profile changed in gene and the differential proportion of transcriptome isoforms of genes with alternative splicing events among horse, donkey and their hybirds, *TNNC2* and *RYR1* genes are my focus because *TNNC2* and *RYR1* genes were significantly enriched in the Muscle contraction pathway and these two genes exhibited dominace from the expression profile of gene in the muscle tissue of mule. Moreover, the proportion of transcriptome isoforms generated from these genes are varied when these genes expressed in the muscle tissue of mule and horse. *TNNC2* gene plays an important role in overcoming the inhibitory effect of troponin complex on actin filament (Townsend et al., 1997; Strausberg et al., 2002). In addition, *TNNC2* gene acts as a calcium release channel in the sarcoplasmic reticulum and a connection between the sarcoplasmic reticulum and the transverse tube(Fujii et al., 1991; Yan et al., 2015). *RYR1* plays a signal role in embryonic skeletal muscle formation. There is a correlation between *RYR1* mediated Ca^2+^ signaling and expression of multiple molecules involved in key myogenic signaling pathways, with less type I muscle fibers when the amount of *RYR1* is reduced (Filipova et al., 2016). Skeletal muscle fibers can be divided into two types: slow contraction type (type I) and fast contraction type (type II)(Talbot and Maves, 2016). Type I muscle fibers contract more effectively over a long period of time and are used primarily for posture maintenance such as head erection or endurance training such as marathon running (Saltin et al., 1977; Yong-Xu et al., 2004). Type II muscle fibers can utilize anaerobic respiration and erupt faster than type I fibers, but they also fatigue more quickly (Zierath and Hawley, 2004).

In brief, our results indicated that alternative splicing events are linked to the heterosis in hybirds. Further experimental verification will be required to reveal the regulatory mechanism of mRNA splicing on gene regulation in hybrids and either of their parents.

## Conclusion

To illuminate the underlying genetic and molecular basis for endurance and cognition heterosis in mule, we conducted comparative analysis of the DEGs and DAS genes between mule and either of their parents and put emphasis on the DEGs and DAS genes which can reflect the heterosis of mule. “Muscle contraction” and “Neuronal System” pathways were significantly enriched by DEGs and DAS genes in the muscle tissue of mule, in which DEGs and DAS genes exhibited over-dominance or dominance effects on muscle endurance and cognition, respectively. Taken together, DEGs and DAS genes can have distinct implications in genetic and molecular mechanisms in heterosis. This study provides valuable resources for future research of muscle endurance echanisms.

## Data Availability Statement

The RNA-seq data from this publication have been submitted to the National Center for Biotechnology Information (https://www.ncbi.nlm.nih.gov/) and assigned the accession SRR4039461, SRR4039465, SRR4039467, SRR4039469, SRR636944, SRR636945, ERR2584210, ERR2584210 and PRJNA387435.

## Author Contributions

S. G. designed the study and conducted bioinformatics analysis.

