## Supplement for "Comparative transcriptome analysis of mule uncovered several heterosis-related genes for improving muscular endurance and cognition"

**Supporting information**

Additional supporting information may be found online in the supporting information tab for this article:

**Table S1** Number of pathways enriched by the DEGs.

**Table S2** Number of pathways enriched by the DAS Genes.

**Figure S1** The PCA of the brain, muscle, and skin tissues in horses, donkeys, mules and hinnies.

**Figure S2** The venn diagrams of differentially expressed genes among three tissues.

**Figure S3** Top 10 pathways enriched in brain tissue by DEGs.

**Figure S4** Top 10 pathways enriched in muscle tissue by DEGs.

**Figure S5** Top 10 pathways enriched in skin tissue by DEGs.

**Figure S6** The venn diagrams of differentially expressed genes and differentially spliced genes in brain, muscle, and skin tissues.

**Figure S7** Proportion of five splicing types of the DAS genes in three tissues.

**Figure S8** Top 10 pathways enriched in brain tissue by DAS genes.

**Figure S9** Top 10 pathways enriched in muscle tissue by DAS genes.

**Figure S10** Top 10 pathways enriched in skin tissue by DAS genes.

**Figure S11** The reads mapping condition of the *TNNC2* among the hinny, mule, donkey, horse in igv.

**Figure S12** The reads mapping condition of the *RYR1* among the hinny, mule, donkey, horse in igv..

**Figure S13** In the comparison between horses and horses, the Pac-Bio full-length transcriptome can cover splicing genes and provable splicing events.

**Table S1** Number of pathways enriched by the DEGs.

| Tissues | Mules vs horses | Mules vs donkeys | Hinnies vs horses | Hinnies vs horses |
|---|---|---|---|---|
| Muscle | 1,030 | 84 | 778 | 979 |
| Brain | 900 | 294 | 665 | 974 |
| Skin | 1,045 | 458 | 798 | 850 |

**Table S2** Number of pathways enriched by the DAS Genes.

| Tissues | Mules vs horses | Mules vs donkeys | Hinnies vs horses | Hinnies vs horses |
|---|---|---|---|---|
| Muscle | 564 | 79 | 503 | 621 |
| Brain | 377 | 108 | 358 | 484 |
| Skin | 289 | 85 | 381 | 341 |

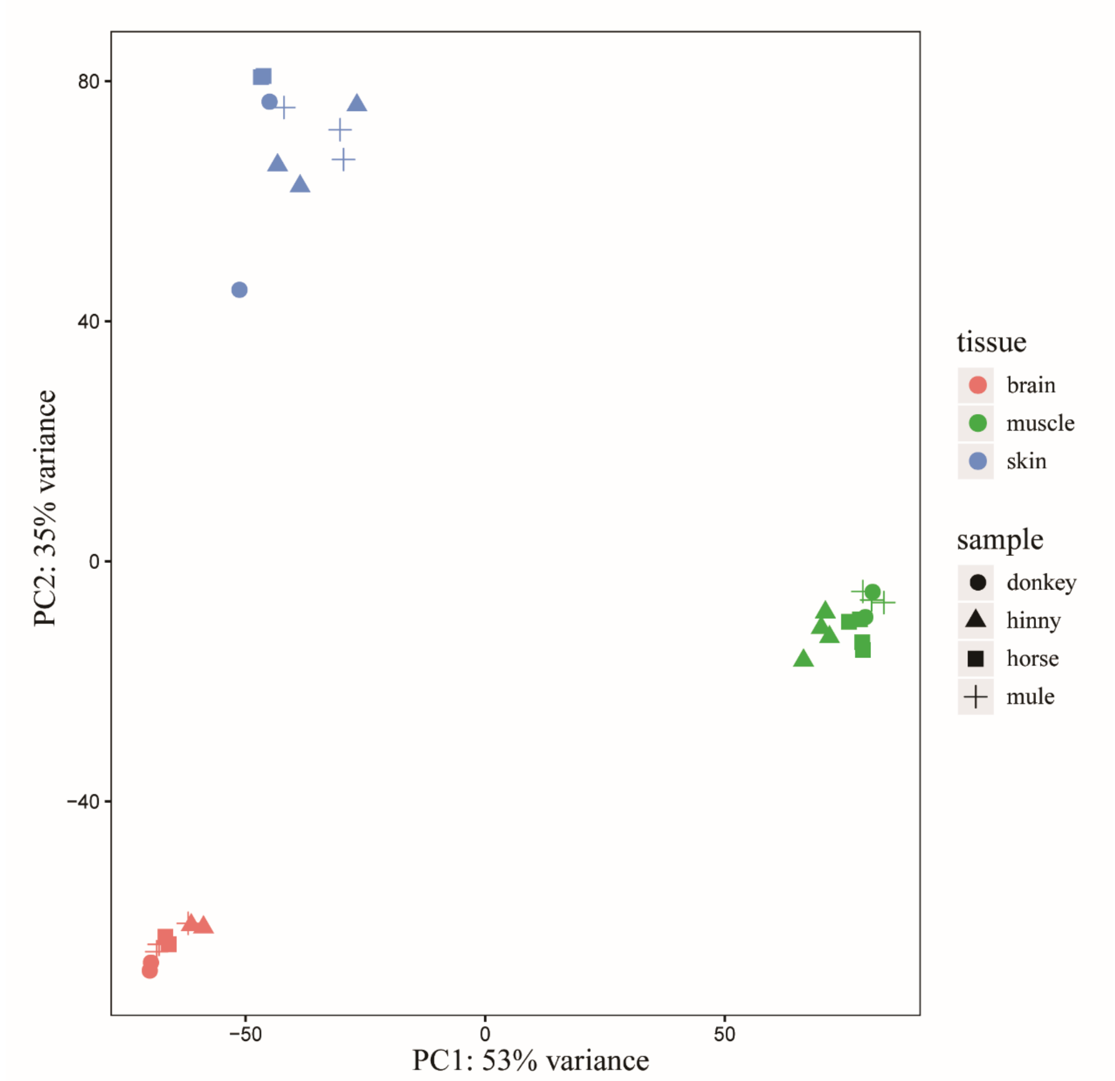


**Figure S1** The PCA of the brain, muscle, and skin tissues in horses, donkeys, mules and hinnies.

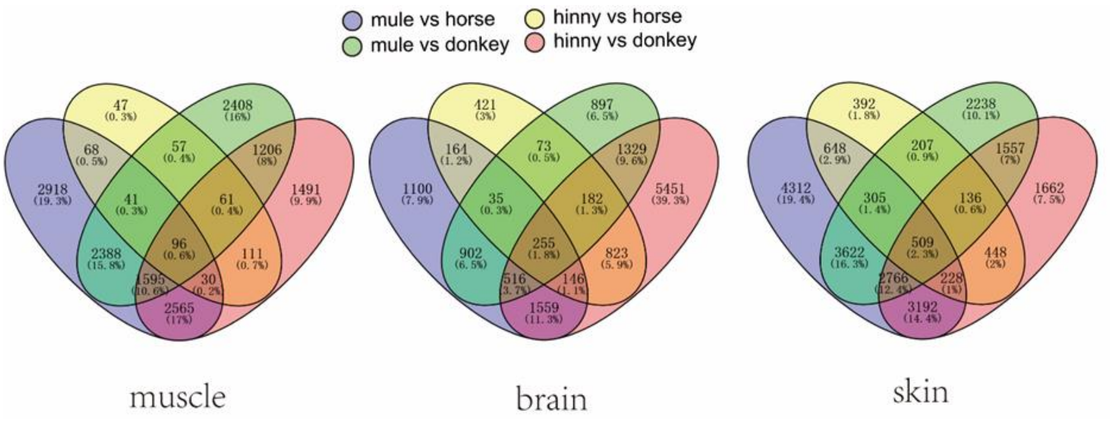


**Figure S2** The venn diagrams of differentially expressed genes among three tissues.

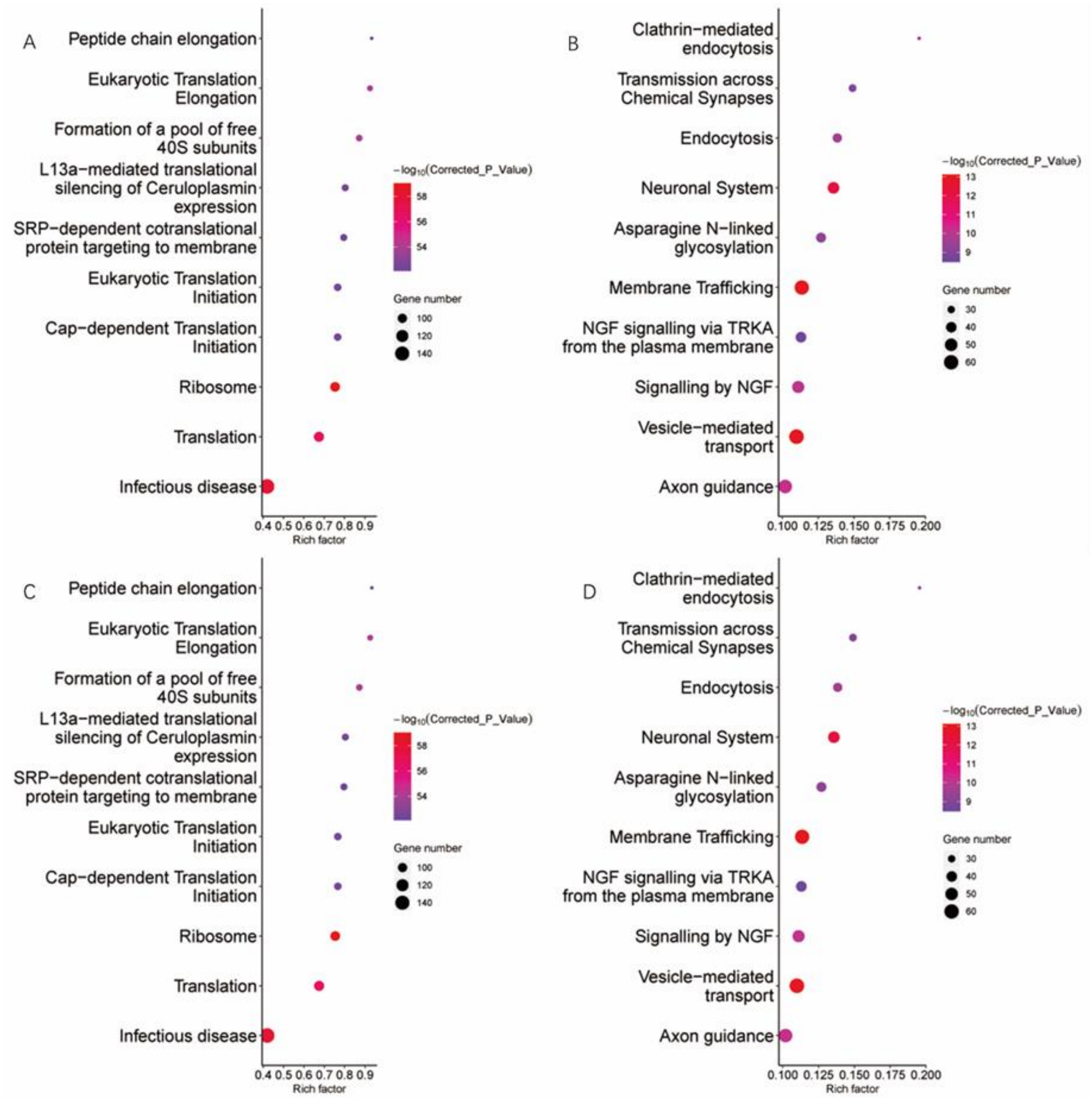


**Figure S3 Top 10 pathways enriched in brain tissue by DEGs**. **A,** pathways enriched by the DEGs between mule vs donkey. **B,** pathways enriched by the DEGs between mule vs horse. **C,** pathways enriched by the DEGs between hinny vs donkey. **D,** pathways enriched by the DEGs between hinny vs horse.

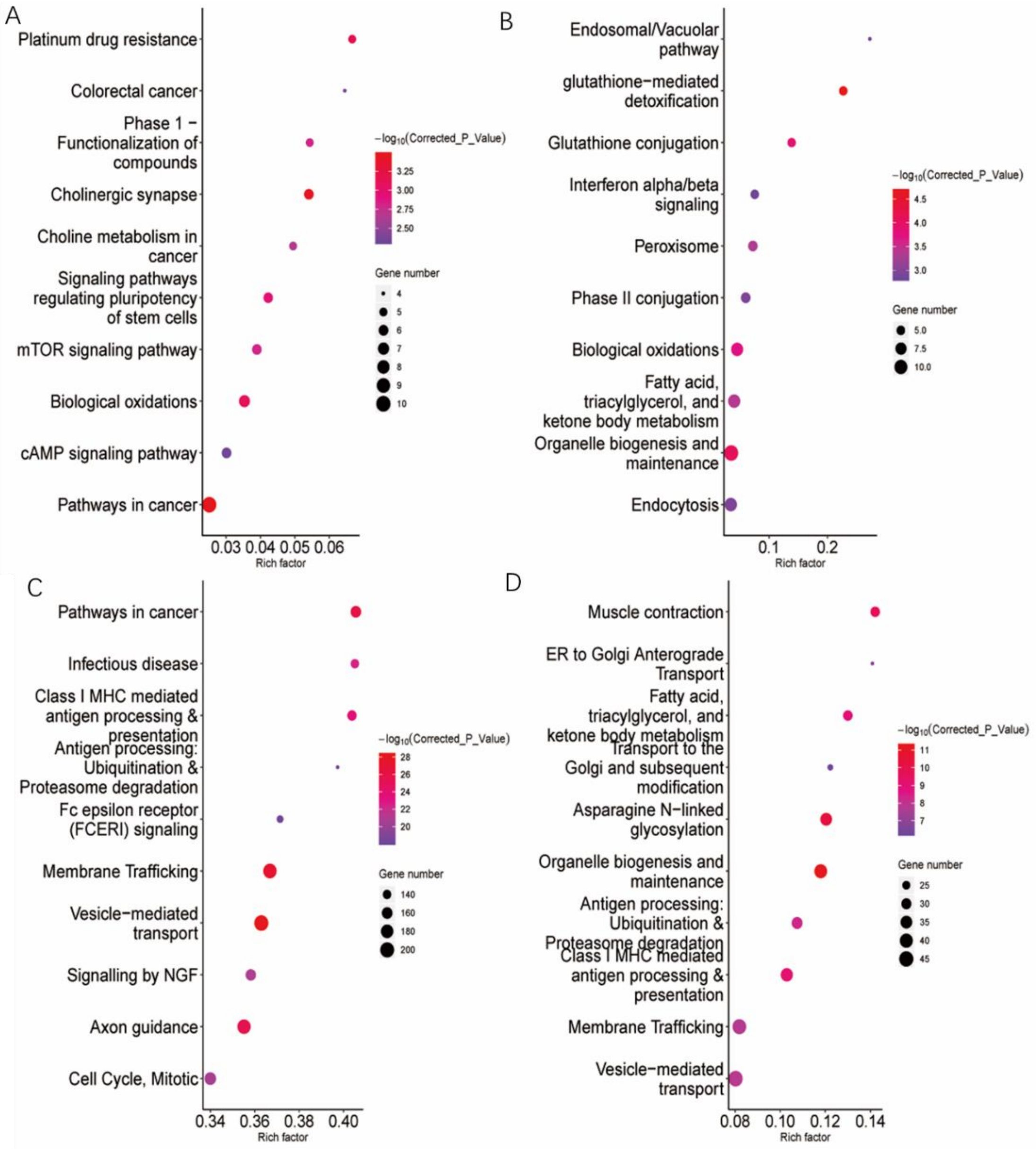


**Figure S4 Top 10 pathways enriched in muscle tissue by DEGs**. **A,** pathways enriched by the DEGs between mule vs donkey. **B,** pathways enriched by the DEGs between mule vs horse. **C,** pathways enriched by the DEGs between hinny vs donkey. **D,** pathways enriched by the DEGs between hinny vs horse.

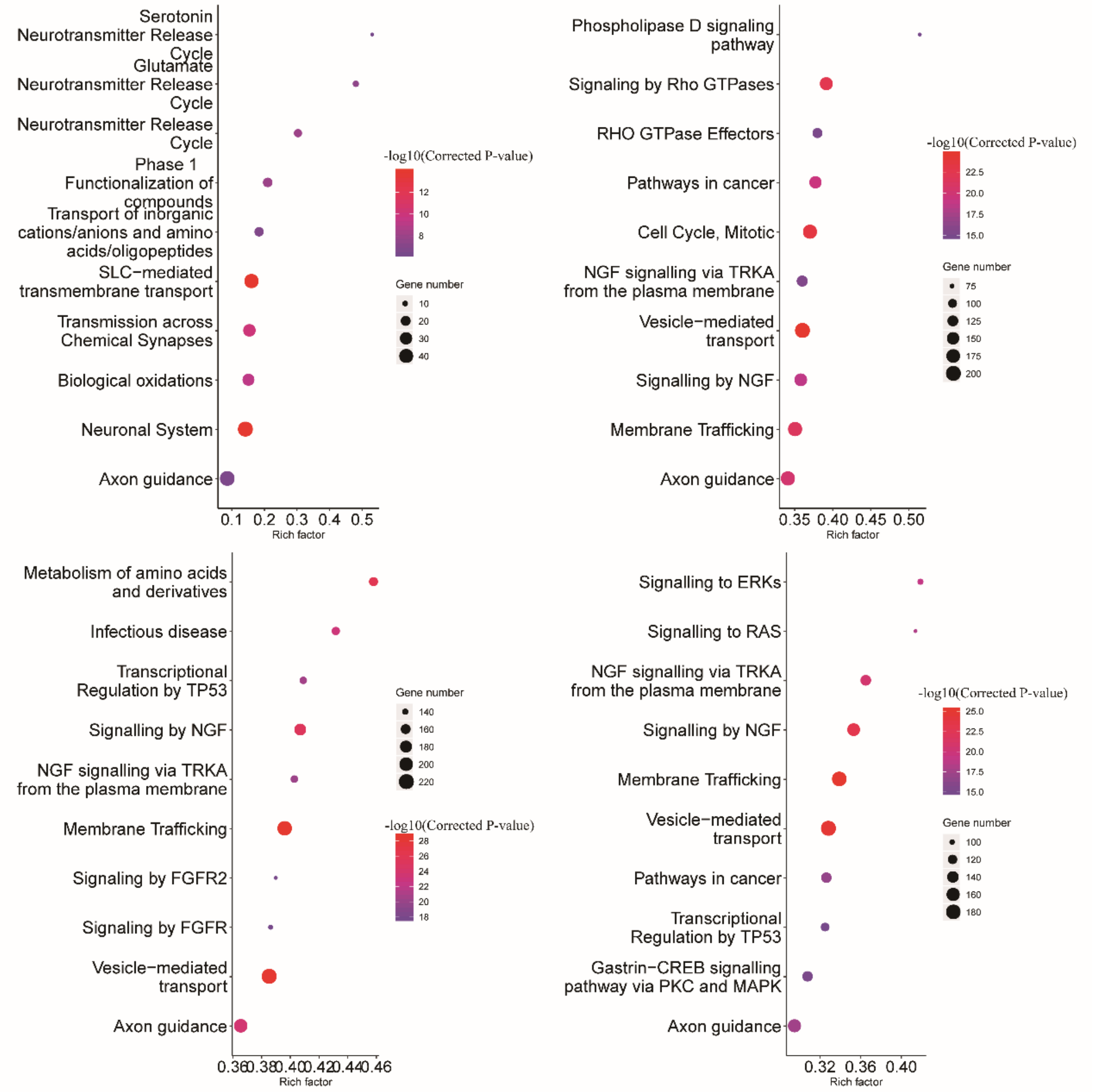


**Figure S5 Top 10 pathways enriched in skin tissue by DEGs**. **A,** pathways enriched by the DEGs between mule vs donkey. **B,** pathways enriched by the DEGs between mule vs horse. **C,** pathways enriched by the DEGs between hinny vs donkey. **D,** pathways enriched by the DEGs between hinny vs horse.

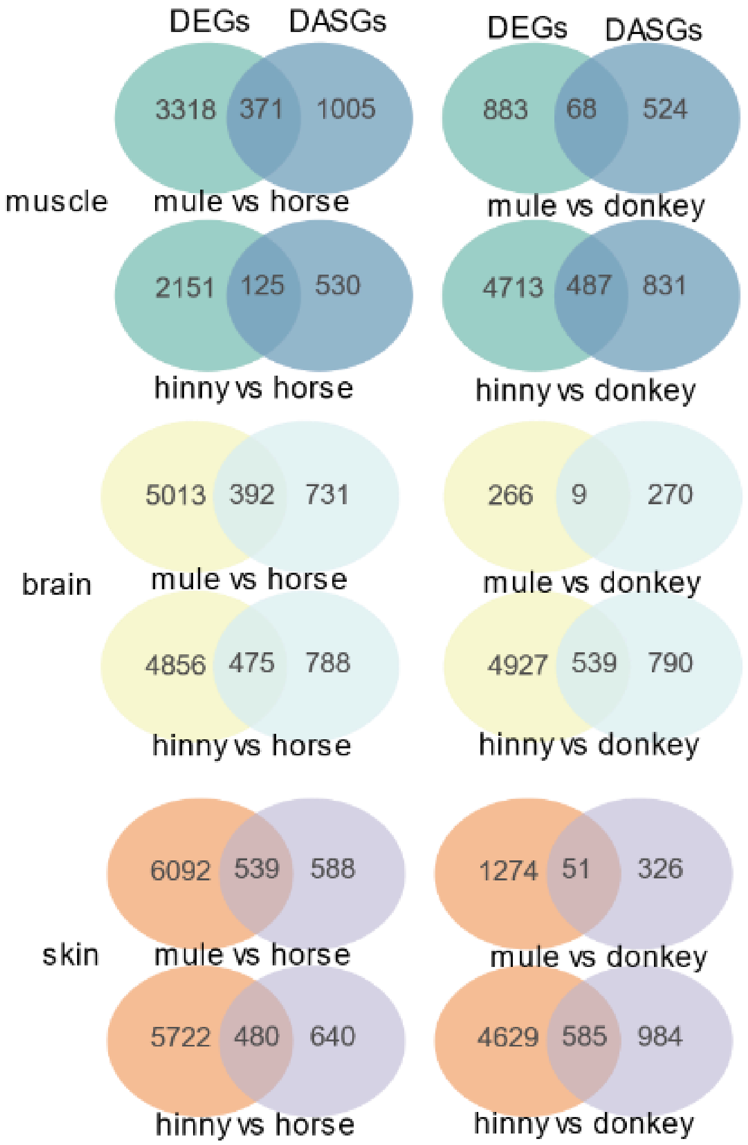


**Figure S6** The venn of differentially expressed genes and differentially spliced genes in brain, muscle, and skin tissues.

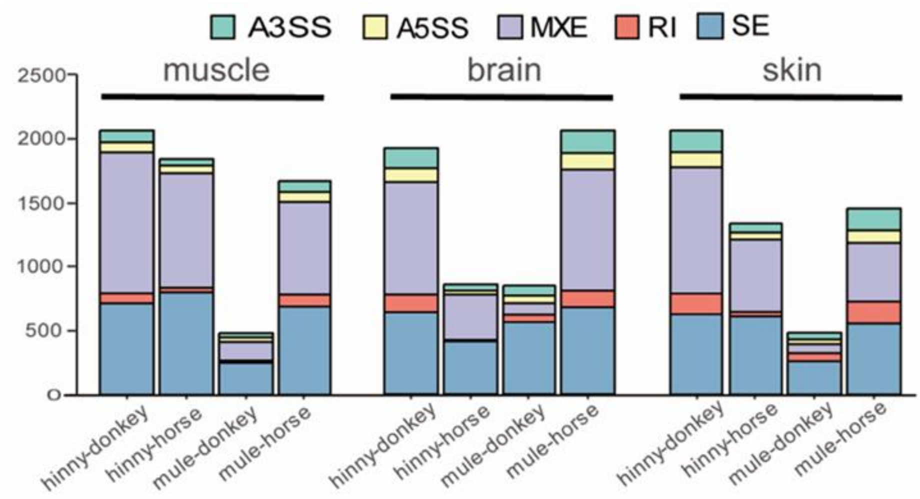


**Figure S7 Proportion of five splicing types of the DAS genes in three tissues. A3SS,** Alternative acceptor site **A5SS,** Alternative donor site **MXE,** Mutually exclusive exons **RI,** Intron retention **SE,** Exon skipping.

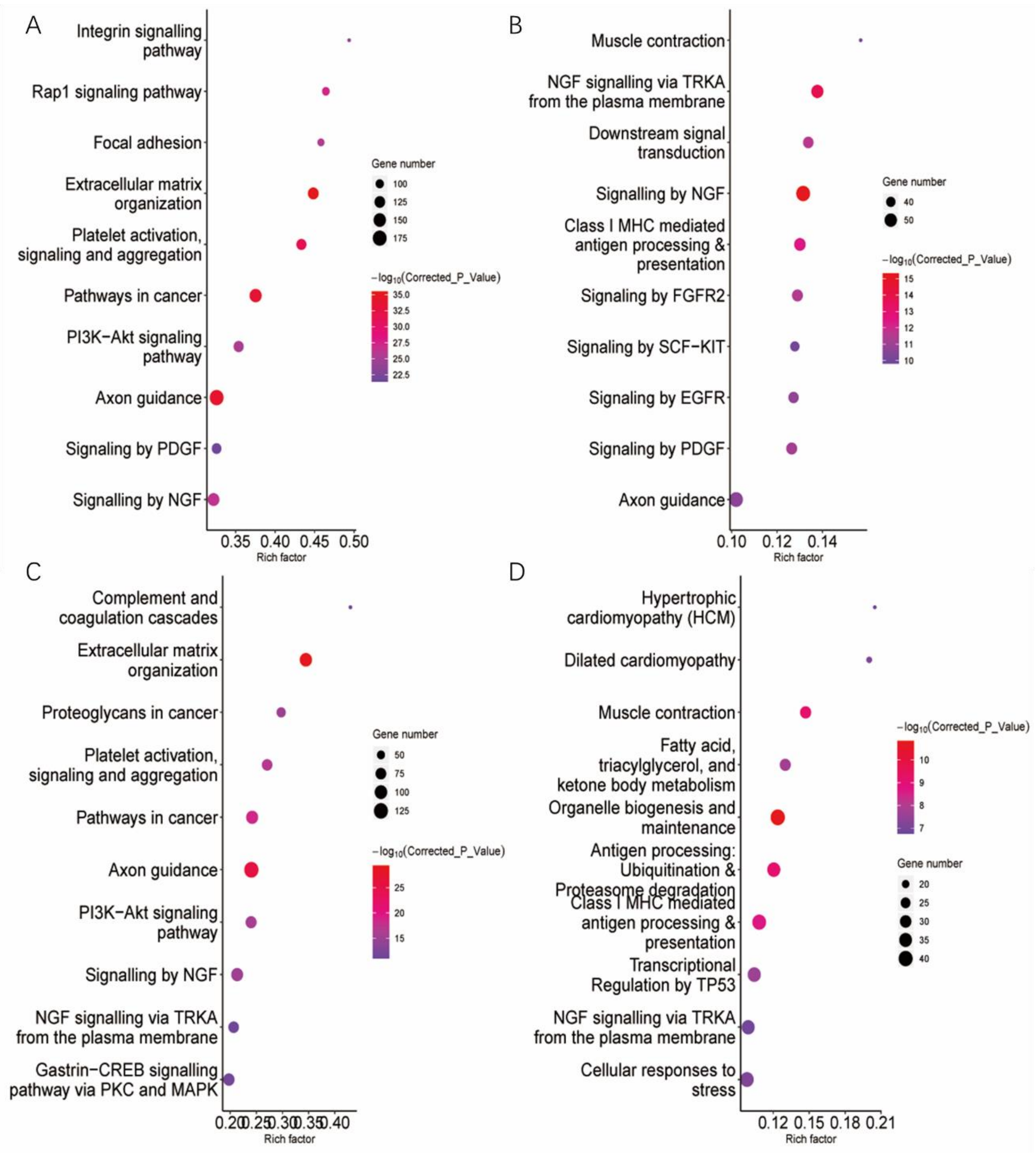


**Figure S8 Top 10 pathways enriched in brain tissue by DAS genes**. **A,** pathways enriched by the DAS genes between mule vs donkey. **B,** pathways enriched by the DAS genes between mule vs horse. **C,** pathways enriched by the DAS genes between hinny vs donkey. **D,** pathways enriched by the DAS genes between hinny vs horse.

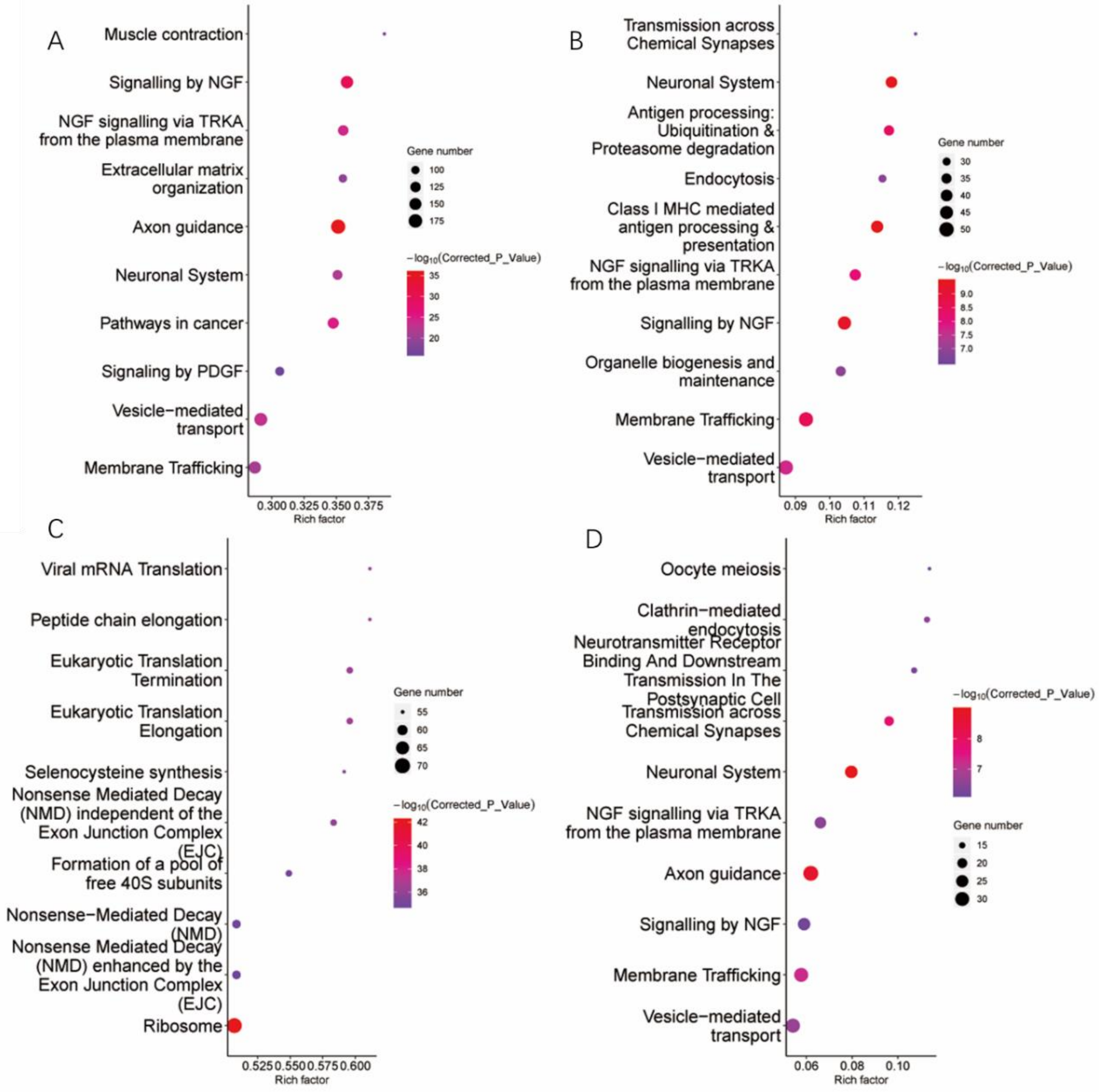


**Figure S9 Top 10 pathways enriched in muscle tissue by DAS genes**. **A,** pathways enriched by the DAS genes between mule vs donkey. **B,** pathways enriched by the DAS genes between mule vs horse. **C,** pathways enriched by the DAS genes between hinny vs donkey. **D,** pathways enriched by the DAS genes between hinny vs horse.

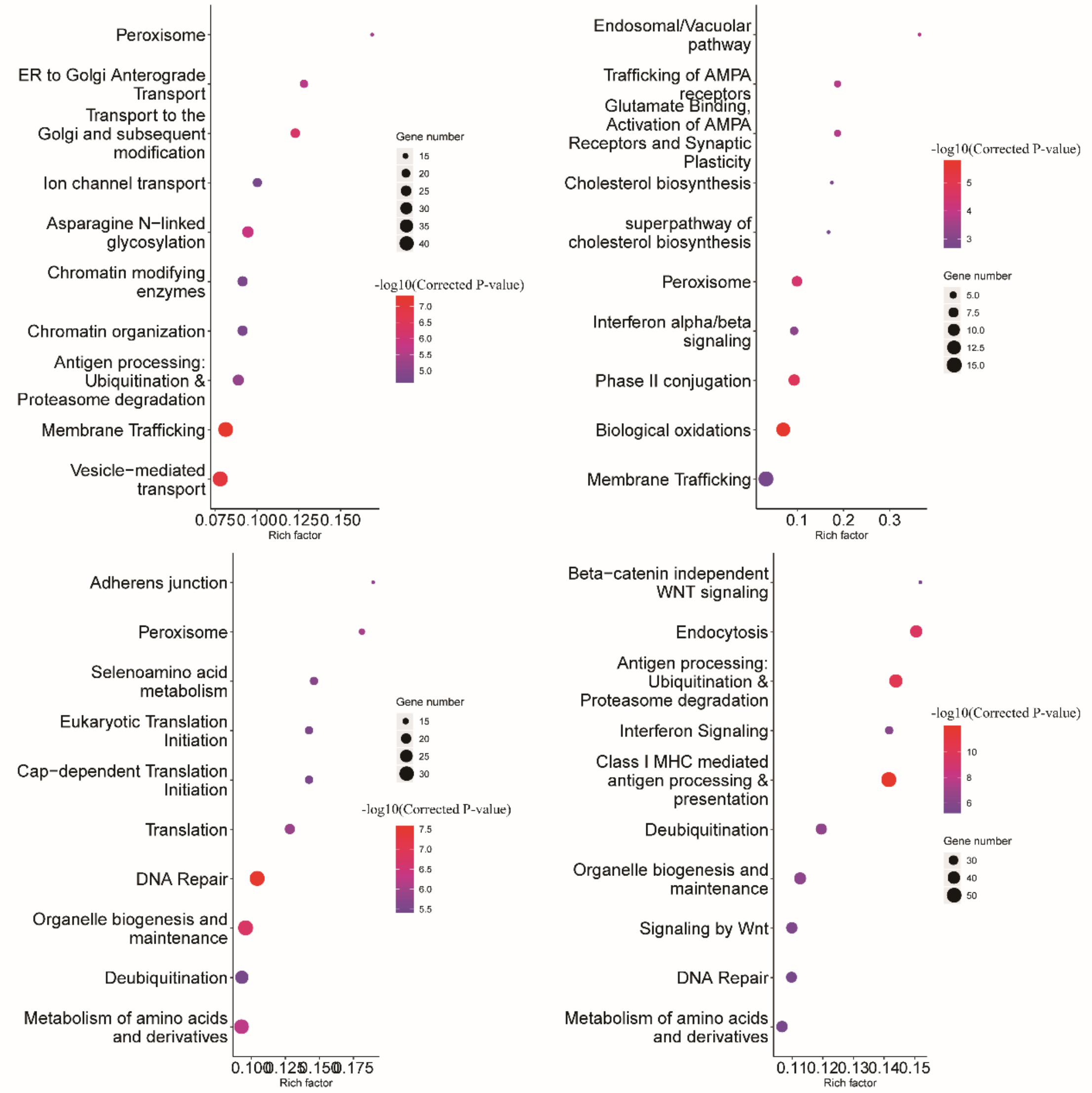


**Figure S10 Top 10 pathways enriched in skin tissue by DAS genes**. **A,** pathways enriched by the DAS genes between mule vs donkey. **B,** pathways enriched by the DAS genes between mule vs horse. **C,** pathways enriched by the DAS genes between hinny vs donkey. **D,** pathways enriched by the DAS genes between hinny vs horse.

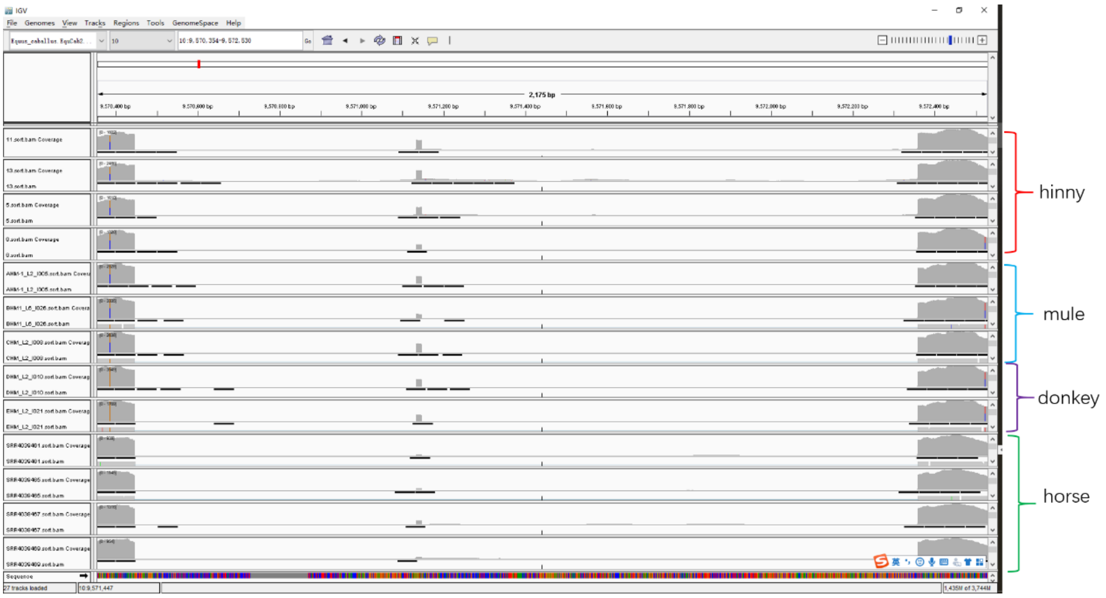


**Figure S11** The reads mapping condition of the *TNNC2* among the hinny, mule, donkey, horse in igv.

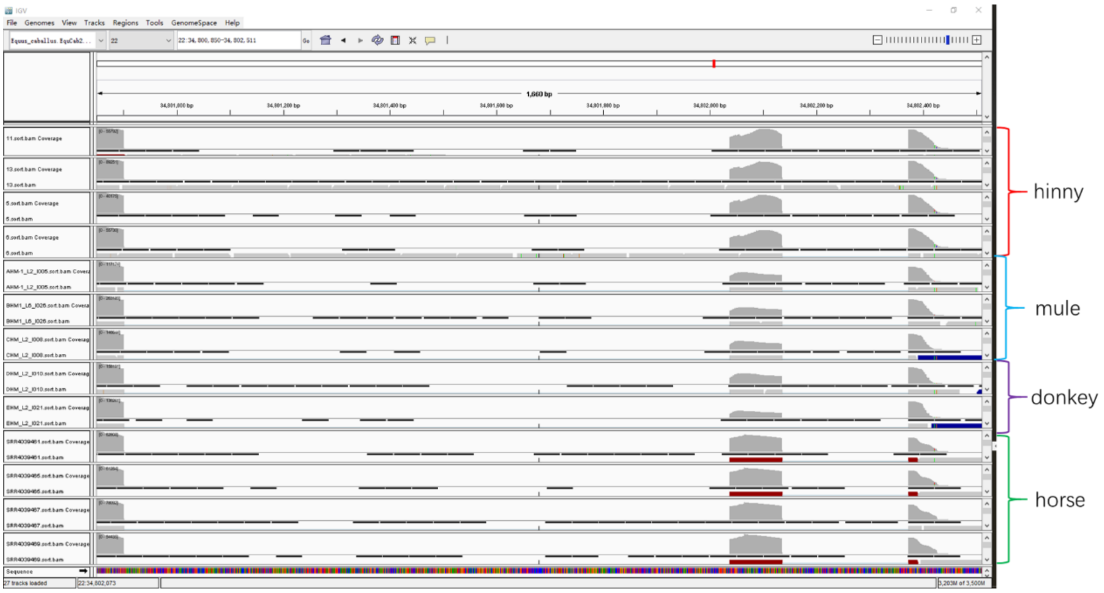


**Figure S12** The reads mapping condition of the *RYR1* among the hinny, mule, donkey, horse in igv.

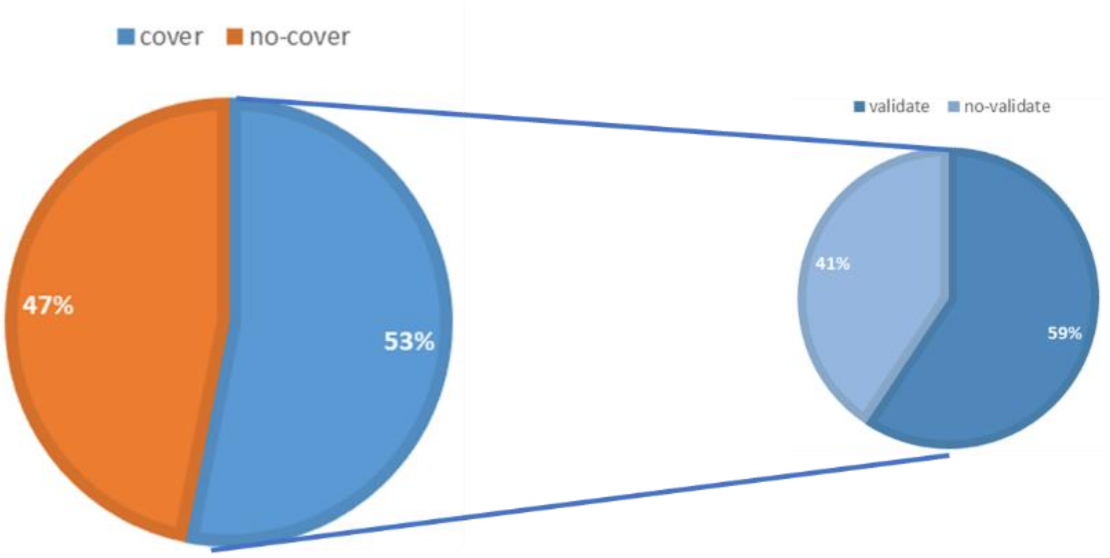


**Figure S13** In the comparison between horses and horses, the Pac-Bio full-length transcriptome can cover splicing genes and provable splicing events.
